# Correlation between antibacterial activity of two *Artemisia* sp. extracts and their plant characteristics

**DOI:** 10.1101/2023.05.30.542989

**Authors:** Abdullah Mashraqi, Mohamed A. Al Abboud, Khatib Sayeed Ismail, Yosra Modafer, Mukul Sharma, A. El-Shabasy

## Abstract

The present study evaluated the potential antibacterial activity of *Artemisia absinthium* L. and *Artemisia herba-alba* Asso. extracts through different organic and aqueous solvents. The tested bacteria were pathogenic types; *Listeria monocytogenes*, *Pseudomonas aeruginosa, Salmonella enterica a*nd *Staphylococcus aureus*. There were different affinities for the studied organic solvents besides aqueous one. The comparative study was accomplished with comparing to the morphological, anatomical and palynological characters. The similarity parameter is obtained. ANOVA test analyzed MIC values for both plant extracts. Pearson Correlation Coefficients were determined for all both plant traits. MIC and MBC values were confirmed on using butanol and diethyl ether extracts besides butanol and chloroform extracts for *Artemisia absinthium* L. and *Artemisia hera alba* Asso against tested pathogenic bacteria respectively as an alternative natural antibacterial inhibitor agent.

## Introduction

Most of secondary metabolites of phytochemicals are derived biosynthetically from phyto-primary metabolites. They can be classified according to their medical values into several groups. The active and medical phytochemicals are alkaloids, tannins, steroids, volatile oils, glycosides, fixed oils, phenols, resins and flavonoids which are efficacy in treating many diseases. These phytochemical constituents are deposited in specific parts such as flowers, leaves, seeds, bark, roots and fruits **(Parekh and Chanda, 2007; Farooq and Hafiz, 2010)**.

Most of medicinal plants have been reported to have antimicrobial activates against many infectious and pathogenic microorganisms. Currently, scientists prefer natural substituents to synthetic additives because they have no side effects besides are safe to the environments. Therefore, many published articles are focused on using plant extracts as natural antibiotics with the recommendations from therapeutic physicians **(Nurul *et al*., 2022; Mariola *et al*., 2022)**. *Artemisia* genus belonging to family Asteraceae, has significant economic importance. It possesses many medical values like antimicrobial, antitumor, antirheumatic and antispasmodic properties. Many species are used as medicinal plants where as others are allergenic and toxic **(Adil *et al*., 2019a; Pedja *et al*., 2019)**. This genus comprises about 500 species presented in the northern hemisphere of the earth as Asia, America and Europe **(Hanan *et al*., 2020)**. The infrageneric taxonomy of this economically important genus is ongoing and ambiguous because the complexity in plant structures. The morphology, anatomy, pollen grain and cypselas of *Artemisia* and its allies are challenging tasks for plant taxonomists. This differentiation may result from different distributions in diverse conditions where *Artemisia* species have a wide range of geographical adaptations in a variety of habitats mainly arid and semiarid regions **(Adil, 2020).**

The objective of this study is focus on using *Artemisia absinthium* L. and *Artemisia herba-alba* Asso. plant extracts with different organic and aqueous solvents against selected pathogenic bacteria. These plant species are belonged to different geographical regions. The antibacterial activity of these plant species is correlated with other plant taxonomical studies as morphological, anatomical and palynological ones.

## Material and Methods

### 1. Plant collection and identification

The plant material *Artemisia absinthium* L. was collected from Faifa mountains, Jazan Province, KSA, which is located at 17°15′55.3′′ latitude, 43°06′47.1′′ longitude, 383 m elevation in September 2022 whereas *Artemisia herba-alba* Asso. was collected from Wadi El-Sheikh area, Sinai, Egypt, which is located at 28°50′15.3′′ latitude, 33°55′30.1′′ longitude, 77 m elevation in July 2022. (**Fig. 1**) (**Map 1**).The identification of studied plant species was issued by the herbarium of the Biology Department, College of Science, Jazan University (JAZUH). The whole plant species were dried in hot air oven at 50°C 24 hr and blended into powder, this powder of plant materials were used for aqueous and other organic solvents; acetone, butanol, chloroform, diethyl ether, ethanol and methanol. Each whole plant extract was prepared by standard methods of **Yasser *et al*. (2019)**. 20 g of each plant material was placed in 250 ml of studied solvent then stirred using a water bath at 45^0^ C for 10 h. After that, the plant residue was separated by filtration through a Whatman No. 1 filter paper and the solvent was evaporated under vacuum on a rotary evaporator at 40^0^ C (Rotavapor R-200, Büchi Labortechnik, Flavil, Switzerland) except for aqueous solvent. The resulting residue was dissolved in 50 ml dimethyl sulfoxide (DMSO) and stored at −20°C for further use. The last step is not preceded during aqueous solvent preparation. Three replicates were performed for each solvent extraction and standard deviations were obtained **(Maria *et al*., 2023)**.

**Fig. 1.**
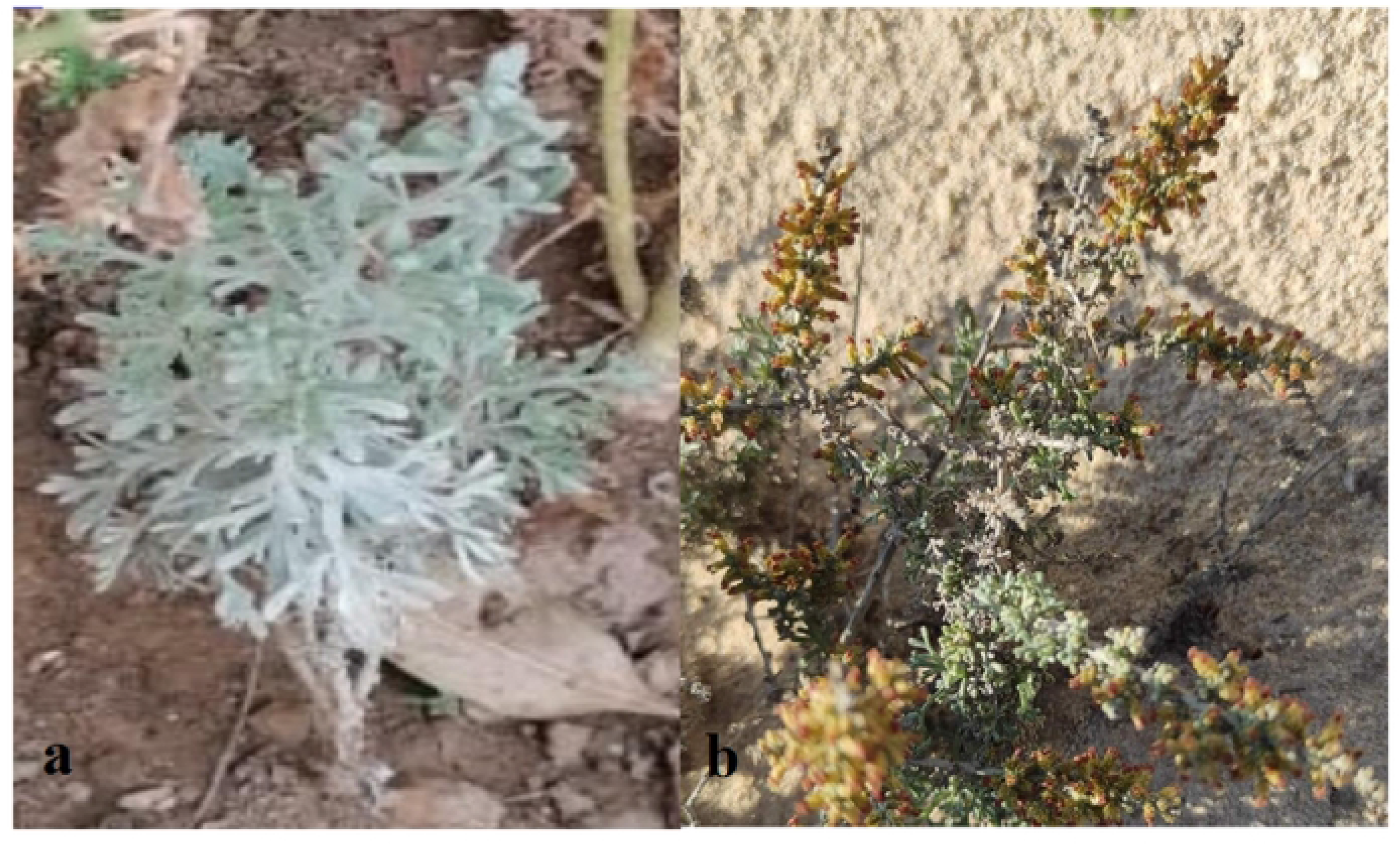
The studied plant species; a. *Artemisia absinthium* L., b. *Artemisia herba-a/ba* Asso.

**Map 1.**
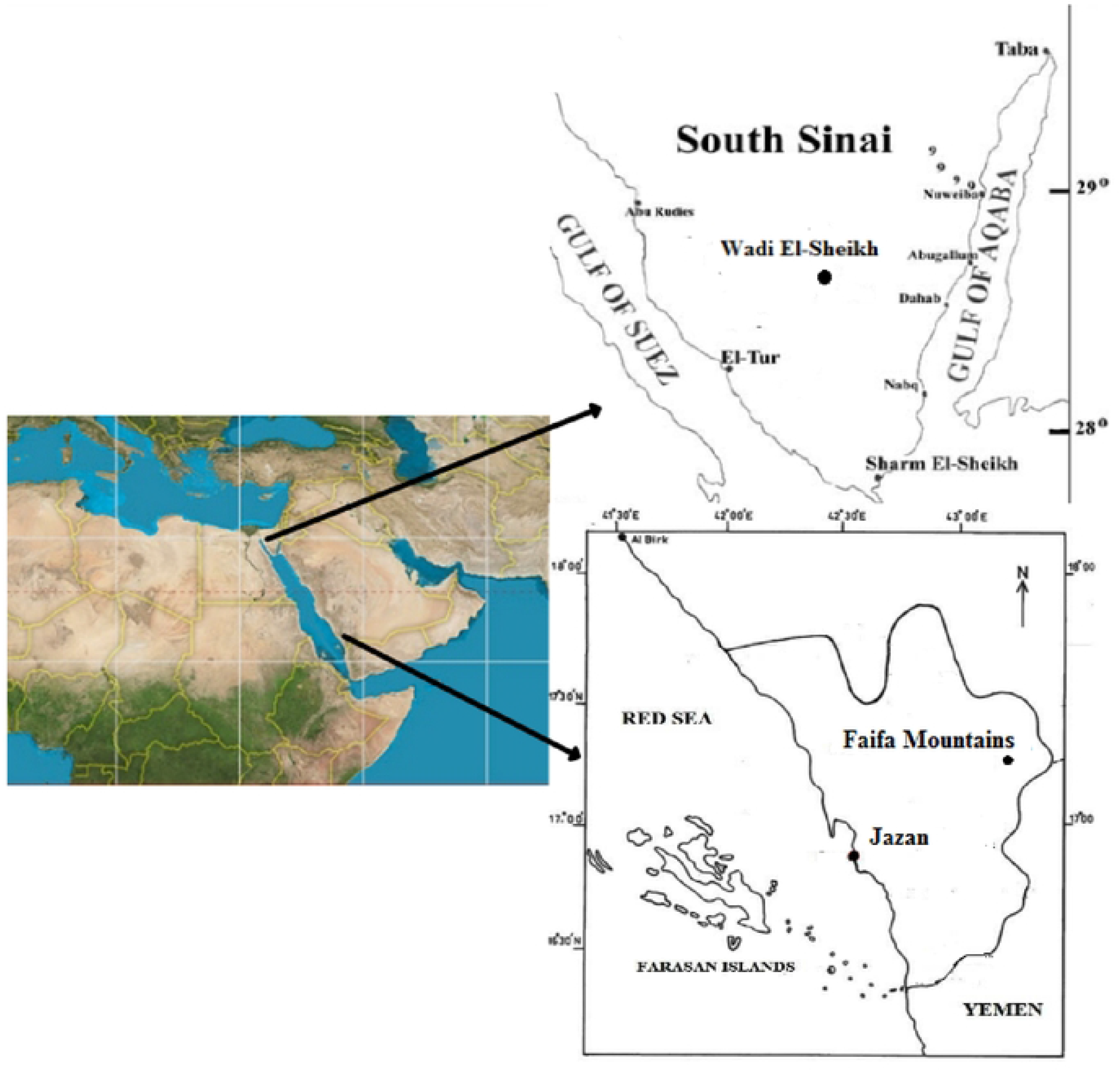
The studied area (E. Atta *et al.,* 2022; Raafat *et al.,*2006).

### 2. Bacterial strains

The antibacterial potency of each plant extract was assessed using four pathogenic bacterial strains. They are isolated, identified and international accredited by ATTC international accredited company.

American Type Culture Collection (ATCC) is established in 1925 when a committee of scientists recognizes the need for a central collection of microorganisms that scientists worldwide can use to conduct their research to advance the science of microbiology. ATCC catalogs published more than 2,000 strains with extensive cross-referencing complete information with trademarks. All summarized data of studied bacterial strains were presented in **(Tables 1, 2)**.

**Table 1.** ATCC clinical database of studied bacterial strains **(Shahid *et al*., 2023).**

| Property | *Listeria monocytogenes* | *Pseudomonas aeruginosa* | *Salmonella enterica* | *Staphylococcus aureus* |
| --- | --- | --- | --- | --- |
| Specific applications | Media testing, Enteric Research, Food testing | Opportunistic pathogen research | Control Culture, Media testing, Preparatory test control, Enteric Research, Emerging infectious disease research, Pharmaceutical and Personal Care, Water testing | CAMP test, Assay of wood, smoke concentrate, Control strain, Control strain for identification, Evaluation of Mueller-Hinton agar, Examination of dairy products, Media testing Quality control strain, Reference material, Susceptibility disc testing, Enteric Research, Water testing |
| Temperature | 37°C | 37°C | 37°C | 37°C |
| Atmosphere | Aerobic | Aerobic | Aerobic | Aerobic |
| Isolation source | Poultry | Clinical isolate from human | Tissue from pools of heart and liver from 4-week-old chickens | Clinical isolate from human |
| Applications | Enteric disease research, Food testing, Infectious disease research, Media testing, Zoonotic disease research | Enteric disease research, Food testing, Infectious disease research, Media testing, Zoonotic disease research | Bioinformatics Enteric disease research Food testing Infectious disease research Media testing Water testing Quality control Pharmaceutical testing | Bioinformatics, Enteric disease research, Infectious disease research, Media testing, Quality control, Water testing |
| Product format | Freeze-dried | Freeze-dried | Freeze-dried | Freeze-dried |
| Storage conditions | 2°C to 8°C | 2°C to 8°C | 2°C to 8°C | 2°C to 8°C |
| Susceptibility profile | - | Gentamicin | - | ampicillin, caryomycin, cephalexin, cephaloglycin, |

**Table 2.** ATCC genomic database of studied bacterial strains **(Shahid *et al*., 2023).**

| Genomic subunits | *Listeria monocytogenes* | *Pseudomonas aeruginosa* | *Salmonella enterica* | *Staphylococcus aureus* |
| --- | --- | --- | --- | --- |
| Number of CDS | 2845 | 6,379 | 4,644 | 2,576 |
| Number of Hypothetical Proteins | 1025 | 2,762 | 1,253 | 928 |
| Number of tRNA | 53 | 4 | 86 | 60 |
| Number of 5s rRNA | 1 | 4 | 8 | 7 |
| Number of 16s rRNA | 1 | 4 | 7 | 6 |
| Number of 23s rRNA | 1 | 4 | 7 | 6 |

*Listeria monocytogenes* **(Murray *et al*.)** Pirie (ATCC® 19111™) is a mesophilic bacterium. It is belonged to Bacillota phylum, bacilli class, caryophanales order, listeriaceae family. It is the causative agent of listeriosis, a Gram-positive, facultatively-intracellular rod capable of causing reproductive disease, neurologic disease and septicaemia in a wide range of hosts. It is regarded as a highly consequential human foodborne pathogen because it is ubiquitous in the environment, as well as it can cause devastating disease in veterinary species, especially ruminants. It is able to tolerate in a wide range of temperature that may replicate when refrigerated at 5°C besides survive in some pasteurization techniques. It can tolerate also in a broad pH range **(Ripolles-Avila *et al*., 2020)**. *Pseudomonas aeruginosa* (Schroeter) Migula (NCTC 10662/ ATCC® 25668™) is an aerobic Gram-negative bacillus opportunistic pathogen. It is able to tolerate in low oxygen conditions because it is regarded as highly versatile bacteria. It can grow in temperature ranging from 4 to 42°C and survive with low level of nutrients. It can withstand on medical equipment and hospital stations where it infects immunocompromised patients. It causes bacteremias, urinary tract infections and pneumonias. It possesses high mortality and morbidity in patients having cystic fibrosis eventually cause respiratory insufficiency then pulmonary damage **(Sara *et al*., 2013)**. *Salmonella enterica subsp. enterica subsp.* (*ex* Kauffmann and Edwards) Le Minor and Popoff serovar Typhimurium (ATCC® 14028™) is classified inside the ɤproteobacteria class and belongs to the *Enterobacteriaceae* family. It is a Gram negative, rod-shaped bacteria and non-spore forming which are facultative anaerobes and mainly show peritrichous motility **(Fábrega and Vila, 2013)**. It is propelled by three or four flagella that arise randomly on its sides and extend to the external medium **(Harshey, 2013)**. *Staphylococcus aureus* subsp. *aureus* Rosenbach strain Seattle 1945 (ATCC® 25923™) is belonged to Bacillota phylum, bacilli class, Bacillales order, Staphylococcaceae family. It is Gram-positive bacteria,, non-spore forming facultative anaerobes and non-motile. It can grow by fermentation or aerobic respiration. It can resist to heat and tolerate to high concentration of salts. It is more virulent opportunistic pathogen that causes a variety of infectious diseases as well as food poisoning **(Harris *et al*., 2002; Todd *et al*., 2014)**.

### 3. Evaluation of the antibacterial activity of plants extract

The disc-diffusion assay was used to determine the antibacterial activity of plant extracts. The media composition for each bacteria strain was illustrated in **(Table 3)**. The steps of culture preparation were carried out by handle procedure. First, Open vial according to enclosed instructions. By using a single tube of #44 broth (5 to 6 mL), withdraw approximately 0.5 to 1.0 mL with a Pasteur or 1.0 mL pipette, the entire pellet was rehydrated. Aseptically transfer this aliquot back into the broth tube and then mix well. Several drops of the suspension were used to inoculate agar slants and plates. Finally the tubes and plate were incubated at 37°C for 24 hours. **(Kesara *et al*., 2023)**. Each plant residue which was dissolved in 50 ml dimethyl sulfoxide (DMSO) dripped on sterile filter paper discs (9 mm diameter, Whatman No. 3 chromatographic paper). Each filter paper discs were loaded with 2 mg of each plant extract. After that, they were placed on agar medium plates inoculated with Reference bacterial American Type Culture Collection strains which were used in antimicrobial screening and incubated at 35°C±2.5°C for 24 h. A paper disc with dimethyl sulfoxide (DMSO) was used as a negative control. Commercial 6-mm diameter discs containing 0.01 mg of Streptomycin were used as a positive control. The diameter of the clear zone surrounding the plant extract loaded discs was measure in millimeter by Vernier caliper as the antibacterial activity of the given extracts **(Mariola *et al*., 2022)**.

**Table 3.** ATCC accredited bacterial media for studied bacterial strains **(Kesara *et al*., 2023)**.

| **ATCC Medium** | ***Listeria monocytogenes*** | ***Pseudomonas aeruginosa*** | ***Salmonella enterica*** | ***Staphylococcus aureus*** |
| --- | --- | --- | --- | --- |
| Medium Name | 44 Brain Heart Infusion Agar/Broth | 3 Nutrient Agar/Broth | 3 Nutrient Agar/Broth | 18 Tryptic Soy Agar/Broth (Soybean-Casein Digest Medium, USP) |
| **Agar Medium Composition** | **52 g Brain Heart Infusion Agar (BD 211065), 1000 ml DI Water** | **23 g Nutrient Agar (BD 213000), 1000 ml DI Water** | **23 g Nutrient Agar (BD 213000), 1000 ml DI Water** | **40 g Tryptic Soy Agar (BD 236950), 1000 ml DI Water** |
| Broth Medium Composition | 37 g Brain Heart Infusion Broth (BD 237500), 1000 ml DI Water | 8 g Nutrient Broth (BD cat 234000), 1000 ml DI Water | 8 g Nutrient Broth (BD cat 234000), 1000 ml DI Water | 30 g Tryptic Soy Broth (BD cat 211825), 1000 ml DI Water |
| **Brain Heart Infusion Composition** | **200 g infusion from Calf Brains, 250 g infusion from Beef Hearts, 10 g Proteose Peptone, 2 g Dextrose, 5 g NaCl, 2.5 g Na<sub>2</sub>HPO<sub>4</sub>, 1000 ml DI Water** | Nil | Nil | Nil |
| Nutrient Agar Composition | Nil | 3 g Beef Extract, 5 g Peptone, 15 g Agar | 3 g Beef Extract, 5 g Peptone, 15 g Agar | Nil |
| **Tryptic Soy Agar Composition** | Nil | Nil | Nil | **17 g Tryptone, 3 g Soytone, 2.5 g Dextrose, 5 g NaCl, 2.5 g K<sub>2</sub>HPO<sub>4</sub>, 15 g Agar** |

### 4. Determination of the minimum inhibitory concentration of plant extracts

MIC (minimum inhibitory concentration) evaluation is the lowest antimicrobial concentration that inhibits microbial growth after 24 h. of incubation. It was determined for each tested bacteria by a serial micro-dilution of plant extract in DMSO solution in the range of 6.36 to 392.0 mg/L. following the protocol described by **Kostíc *et al*. (2017)**. The inhibition zones were estimated by Vernier caliper and documented against the concentrations of the effective plant extracts.

### 5. Assessment of the Minimum Bactericidal Concentration (MBC)

MBC (minimum bacterial concentration) is the lowest plant extract concentration that showed no bacterial growth after MIC approach. The freshly inoculated agar plates were incubated at 37 °C for 24 h. For each plate, three separate biological replicates were performed, and no colonies on plates were recognized **Joshi (2010)**.

### 6. Plant characterizations

Plant characterizations were determined as qualitative and quantitative characters. Qualitative characters are descriptive plant parts while quantitative ones are measurable. Morphological characters are represented in major seven groups; habitat, leaf, stem, petiole, bract, inflorescence, Cypselas. Each one comprised many sub-characters which denoted with (+) or (-). Moreover, anatomical characters were represented in major five groups; leaf, stem, root, leaf epidermis and stomata. All anatomical ones were observed and measured according to **(Muhammad *et al*., 2010; Pedja *et al*., 2019; Shmygareva *et al*., 2019)**. Cypsela features were refered to (**Rubina and Qaiser 2008)**. (+) refers to present while (-) refers to absent. SEM photos for pollen grains were captured according to **(Adil *et al*., 2019b)**. Polar axis, equatorial axis, P/E sphericity, exine thickness and aperture length were measured by (**Muhammad *et al.,* 2010; Adil *et al*., 2019b).** All other palynological characters were measured from the previous photos as new investigations. They were classified also as qualitative and quantitative characters. Glossary for all abbreviations was presented in the supplementary section.

### 7. Statistical Analysis

Pearson correlation coefficients among MIC values among Minimum Inhibitory Concentration (MIC) of *A. absinthium* and *A. herba alba* extracts against the tested bacterial strains was achieved according to **(Shaban, 2005 and Tamhane, 2009)**. P values for significance tests based on degrees of freedom were determined according to **Dutilleul (1993)** approach. MIC values for each plant extract were subjected to statistical analysis to work out ANOVA to compare means. Standard error was calculated following **T. Al faifi and A. El-Shabasy (2021)**. Statistical tests were conducted using SPSS software (ver. 22) for Windows **(E. Atta *et al*., 2022)**. The representations of (SLR) of the significant relationships were followed **(Maindonald, 1992; Miller and Franklin, 2002)**. They were found for each bacterial strain against two studied plant extracts from one side and for comparative data for both plant species from other side. They were done by the linear regression approaches which explored the extent effect for comparative parameters.

## Results

### 1. Antibacterial analysis

The antibacterial activities of different solvent extracts of the two studied plant species in terms of the diameters of the inhibition zones (IZ), the minimum inhibitory concentrations (MIC) and Minimum Bacterial Concentration (MBC) were presented in **(Tables 4-9) (Fig. 2-7)** referred with alphabetical letters written as abbreviations on bacterial cultures.

**Fig. 2.**
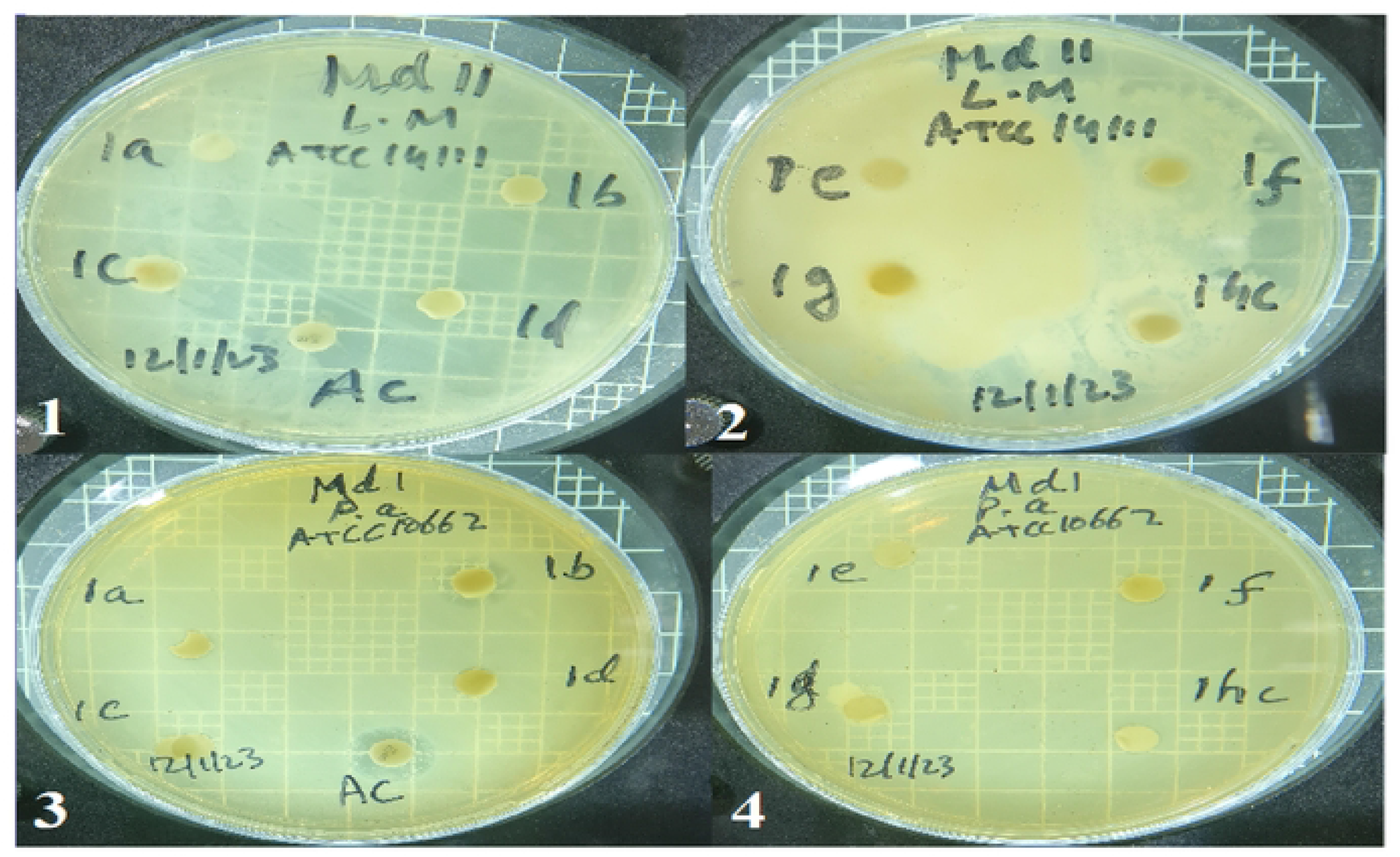
Diameter of Inhibition Zone (IZ) of *A. absinthium* against; 1-2. *Listeria monocytogenes* (L m) and 3-4. *Pseudomonas aeruginosa* (P a) by using different extract solvents.

**Fig. 3.**
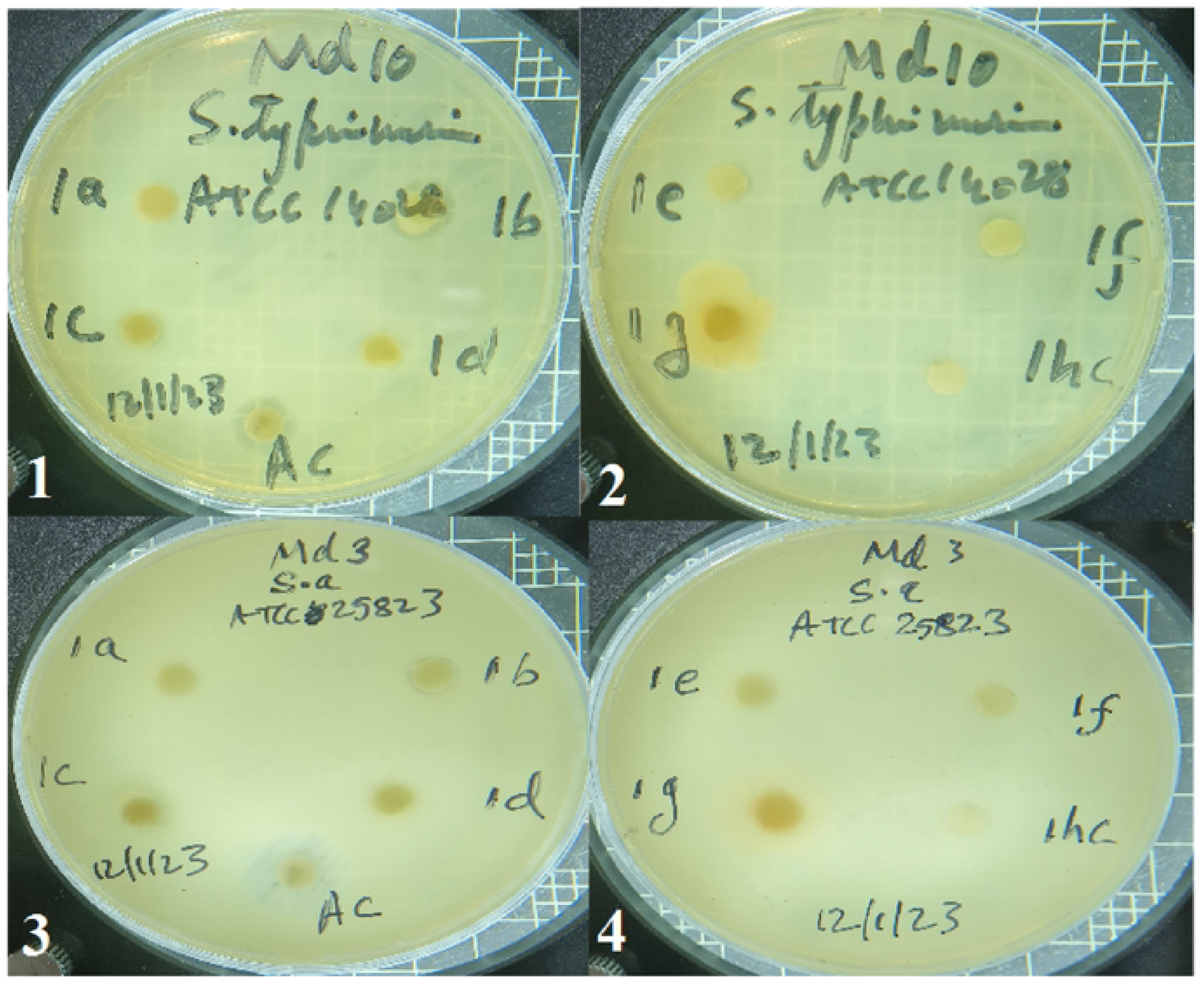
Diameter of Inhibition Zone (IZ) of *A. absinthium* against; 1-2. *Salmonella enterica* (Se) and 3-4. *Sraphylococc11s aureus* (Sa) by using different extract solvents.

**Fig. 4.**
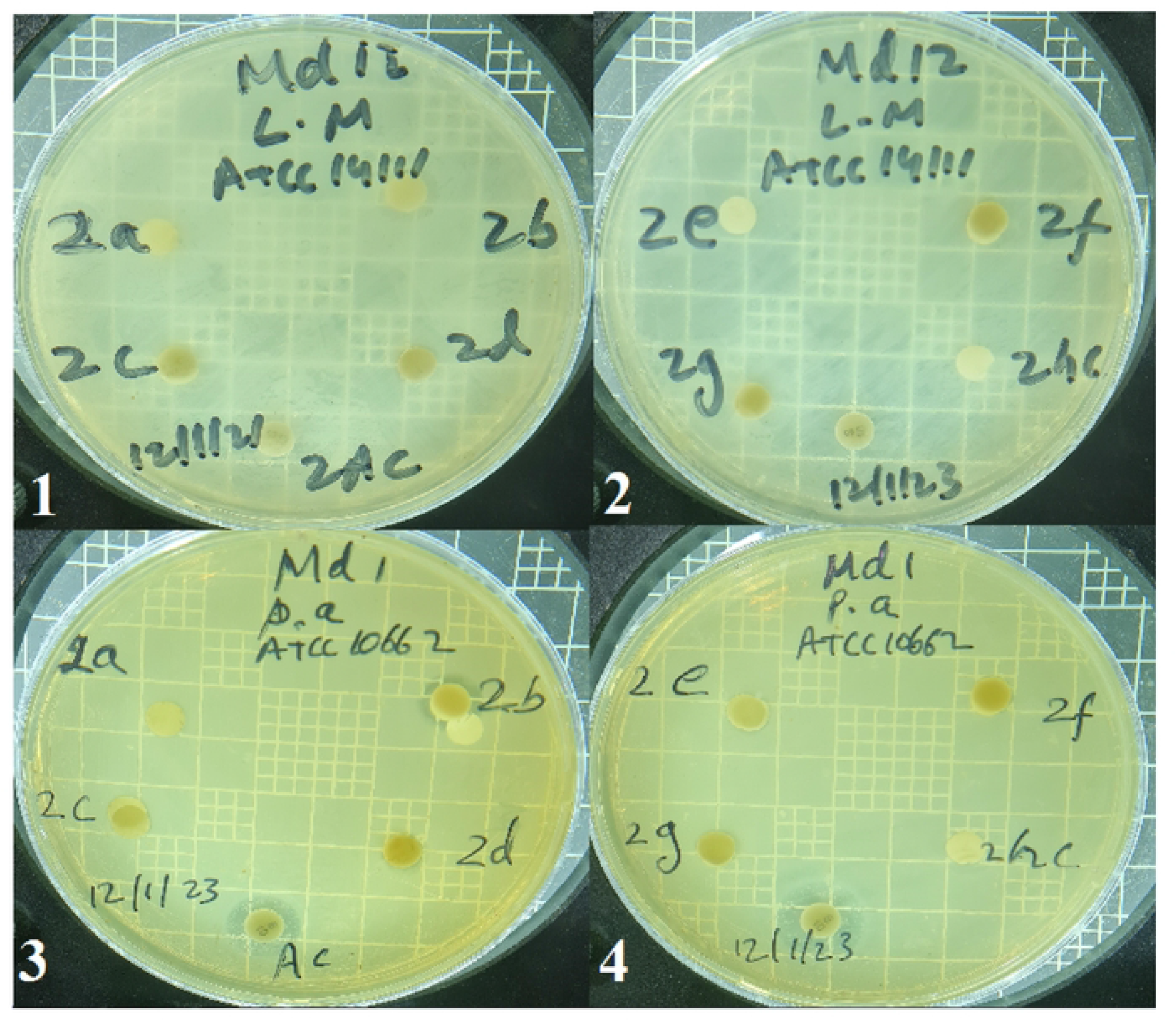
Diameter of Inhibition Zone (IZ) of *A. herba alba* against; 1-2. *Listeria monocytogenes* (L m) and 3-4. *Pseudomonas aeruginosa* (P a) by using different extract solvents.

**Fig. 5.**
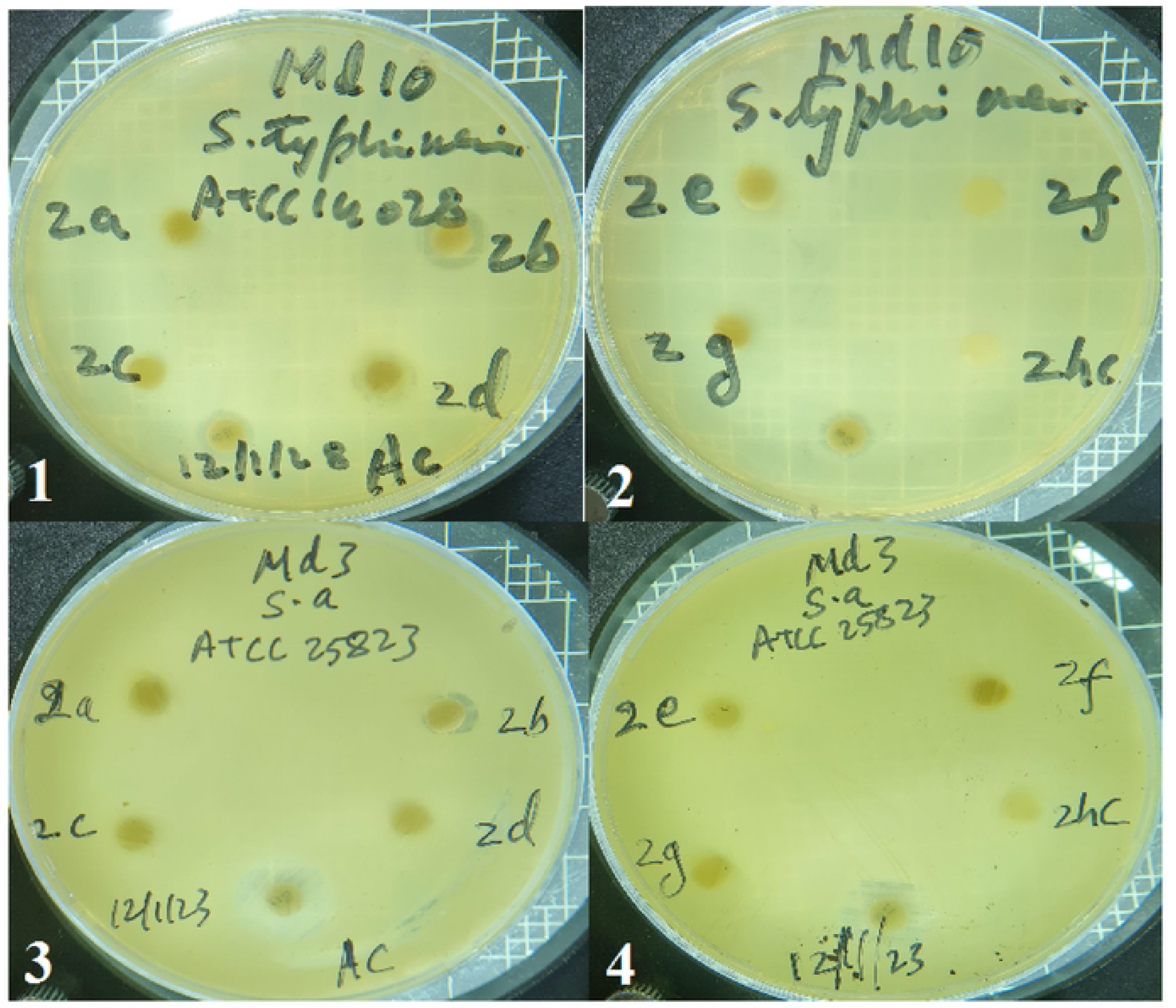
Diameter of Inhibition Zone (IZ) of *A. herba alba* against; 1-2. *Salmonella enterica* (S e) and 3-4. *Staphylococcus aureus* (Sa) by using different extract solvents.

**Fig. 6.**
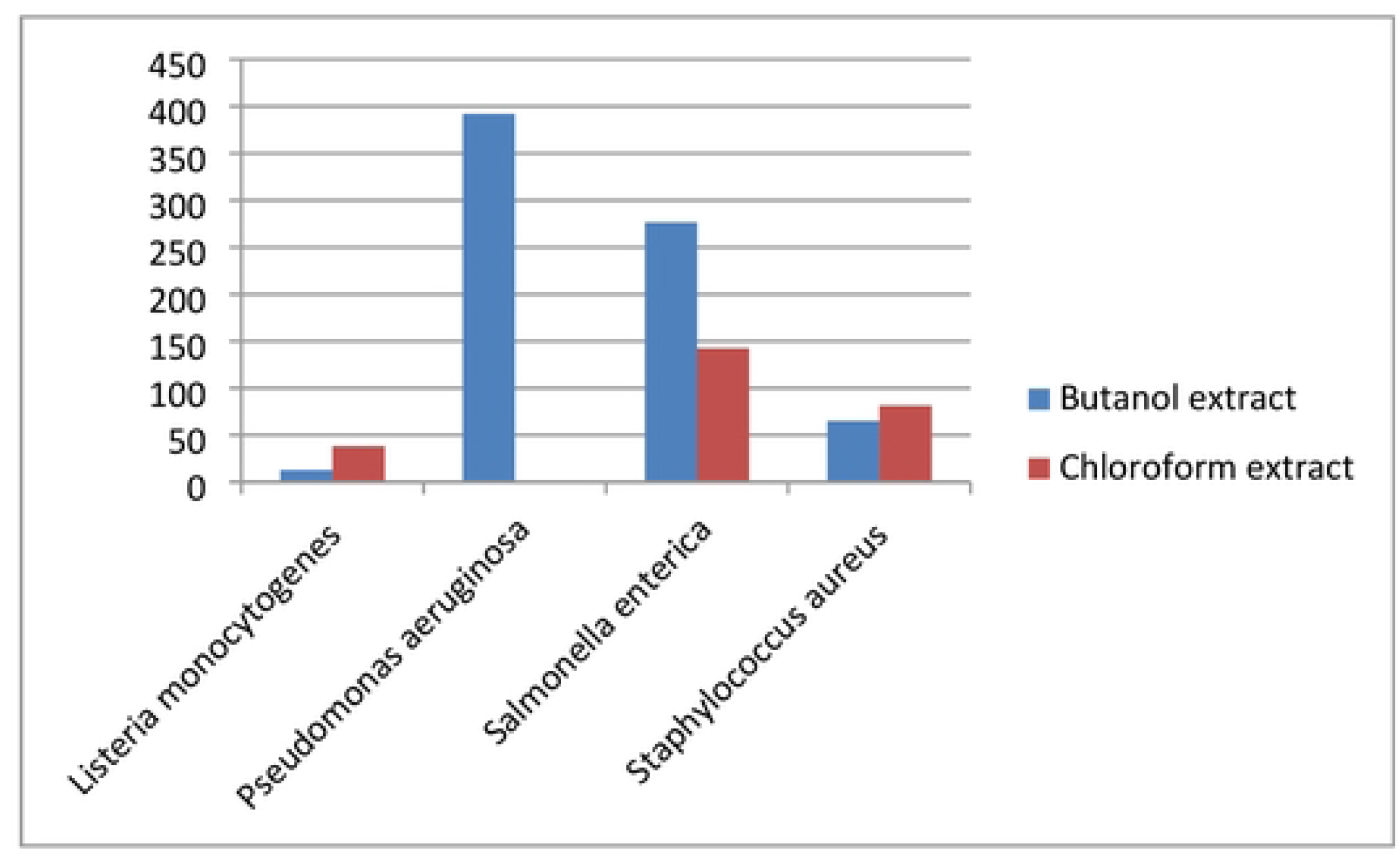
MIC of *A. absinthium* extracts against the tested bacterial strains.

**Fig. 7.**
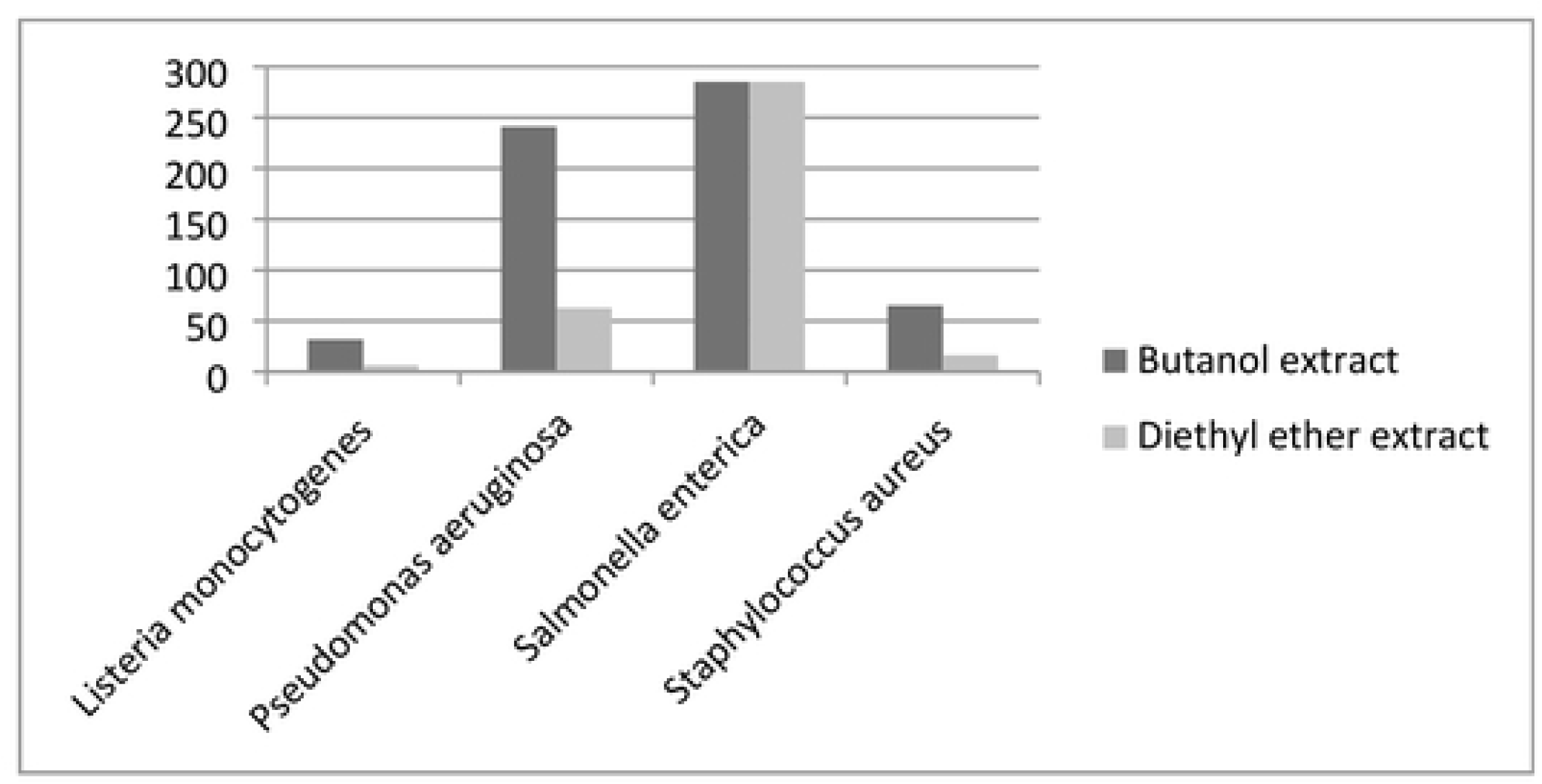
MIC of *A. herba alba* extracts against the tested bacterial strains.

**Table 4.** Antibacterial activity of *A. absinthium* extracts againsts the tested bacterial strains

| Bacterial strain | Diameter of Inhibition Zone (IZ) in mm |  |  |  |  |  |  |  |  |
| --- | --- | --- | --- | --- | --- | --- | --- | --- | --- |
|  | Organic solvents |  |  |  |  |  | Aqueous solvent (g) | DMSO (Negative control) (h) | Streptomycin (Positive control) (AC) |
|  | Acetone (a) | Butanol (b) | Chloroform (c) | Diethyl ether (d) | Ethanol (e) | Methanol (f) |  |  |  |
| *Listeria monocytogenes* | - | 1.0±0.01 | 3.0±0.01 | - | - | 2.5±0.01 | - | - | 12.0±0.01 |
| *Pseudomonas aeruginosa* | - | 5.5±0.58 | - | - | - | - | -2.75±1.71 | - | 7.0±0.01 |
| *Salmonella enterica* | - | 3.88±0.25 | 2.0±0.01 | - | - | - | -11.0±4.08 | - | 9.5±0.01 |
| *Staphylococcus aureus* | - | 2.0±0.01 | 2.5±0.01 | - | - | - | -3.5±0.01 | - | 11.0±0.01 |

**Table 5.** Antibacterial activity of *A. herba alba* extracts against the tested bacterial strains

| Bacterial strain | Diameter of Inhibition Zone (IZ) in mm |  |  |  |  |  |  |  |  |
| --- | --- | --- | --- | --- | --- | --- | --- | --- | --- |
|  | Organic solvents |  |  |  |  |  | Aqueous solvent (g) | DMSO (Negative control) (h) | Streptomycin (Positive control) (AC) |
|  | Acetone (a) | Butanol (b) | Chloroform (c) | Diethyl ether (d) | Ethanol (e) | Methanol (f) |  |  |  |
| *Listeria monocytogenes* | - | 2.5±0.01 | - | 0.5±0.01 | - | - | - | - | 9.0±0.01 |
| *Pseudomonas aeruginosa* | - | 3.38±0.48 | - | 0.88±0.25 | - | - | - | - | 8.0±0.01 |
| *Salmonella enterica* | - | 4.0±0.01 | - | 4.0±0.01 | - | - | - | - | 3.5±0.01 |
| *Staphylococcus aureus* | - | 2.0±0.01 | - | 0.5±0.01 | - | - | - | - | 8.0±0.01 |

**Table 6.**
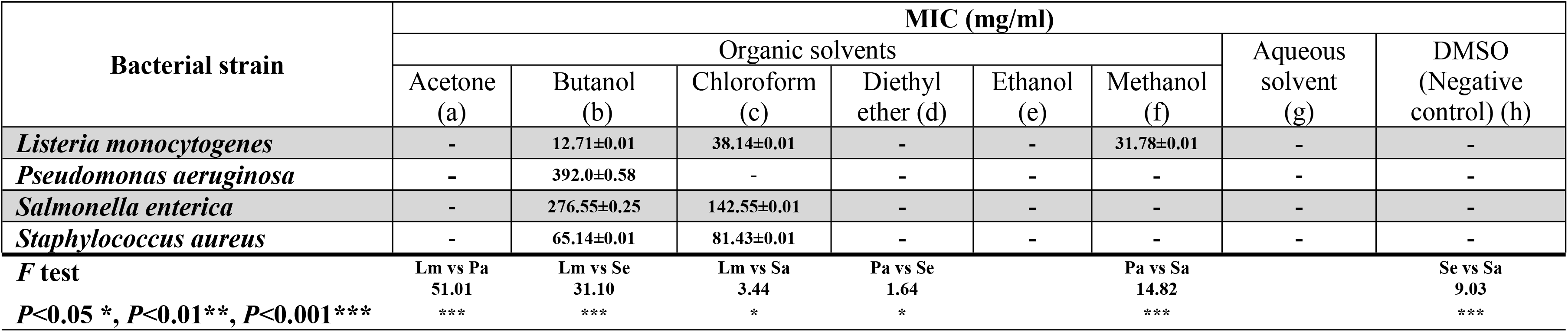
MIC of *A. absinthium* extracts against the tested bacterial strains.

**Table 7.**
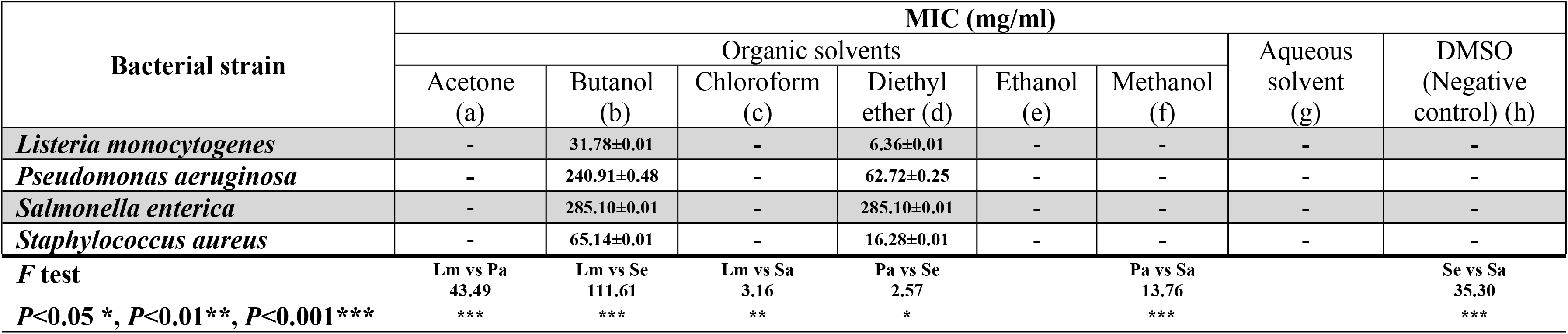
MIC of *A. herba alba* extracts against the tested bacterial strains.

**Table 8.** MBC of the tested bacterial strains against *A. absinthium* extracts.

| Bacterial strain | MBC (mg/ml) |  |  |  |  |  |  |  |
| --- | --- | --- | --- | --- | --- | --- | --- | --- |
|  | Organic solvents |  |  |  |  |  | Aqueous solvent (g) | DMSO (Negative control) (h) |
|  | Acetone (a) | Butanol (b) | Chloroform (c) | Diethyl ether (d) | Ethanol (e) | Methanol (f) |  |  |
| *Listeria monocytogenes* | - | 8.0±0.01 | 24.01±0.01 | - | - | 20.0±0.01 | - | - |
| *Pseudomonas aeruginosa* | - | 2.0±0.58 | - | - | - | - | - | - |
| *Salmonella enterica* | - | 2.0±0.25 | 1.03±0.01 | - | - | - | - | - |
| *Staphylococcus aureus* | - | 4±0.01 | 5±0.01 | - | - | - | - | - |

**Table 9.** MBC of the tested bacterial strains against *A. herba alba* extracts

| Bacterial strain | MBC (mg/ml) |  |  |  |  |  |  |  |
| --- | --- | --- | --- | --- | --- | --- | --- | --- |
|  | Organic solvents |  |  |  |  |  | Aqueous solvent (g) | DMSO (Negative control) (h) |
|  | Acetone (a) | Butanol (b) | Chloroform (c) | Diethyl ether (d) | Ethanol (e) | Methanol (f) |  |  |
| *Listeria monocytogenes* | - | 20.0±0.01 | - | 4.0±0.01 | - | - | - | - |
| *Pseudomonas aeruginosa* | - | 1.23±0.48 | - | 0.32±0.25 | - | - | - | - |
| *Salmonella enterica* | - | 2.06±0.01 | - | 2.06±0.01 | - | - | - | - |
| *Staphylococcus aureus* | - | 4.0±0.01 | - | 1.0±0.01 | - | - | - | - |

Butanol extract of *A. absinthium* revealed antibacterial effects on all tested bacteria. The highest antibacterial activity was exhibited on *Pseudomonas aeruginosa* (IZ 5.5 mm) while the lowest one was on *Listeria monocytogenes* (IZ 1.0±0.01). On the other hand, chloroform extract showed the same effect except on *Pseudomonas aeruginosa*. The highest antibacterial activity was exhibited on *Listeria monocytogenes* (IZ 3.0±0.01 mm) while the lowest one was on *Salmonella enterica* (IZ 2.0±0.01 mm). On contrast, the methanol extract had the only antibacterial activity on *Listeria monocytogenes* (IZ 2.5±0.01 mm). Butanol and diethyl ether of *A. herba alba* extracts had only the antibacterial activity on all tested pathogenic bacteria. *Salmonella enterica* exhibited the highest antibacterial activity on both plant extracts with value IZ: 4.0±0.01 mm while *Staphylococcus aureus* showed the lowest one with values IZ: 2.0±0.01 and 0.5±0.01 mm in butanol and diethyl ether extracts respectively. Moreover, *Listeria monocytogenes* showed the same activity like *Staphylococcus aureus* by using diethyl ether extract.

Despite of aqueous solvents hadn’t any effects on bacterial growth by using *A. herba alba* extracts but there was promoting in growth when *A. absinthium* is used against all tested pathogenic bacteria except *Listeria monocytogenes*. DMSO (negative control) had never any such inhibited activity. Streptomycin (positive control) showed predominantly antibacterial activity against all tested bacteria.

Considering MIC values, *Listeria monocytogenes* showed the lowest value (12.71±0.01 mg/ml) while *Pseudomonas aeruginosa* showed the highest one (392.0±0.58 mg/ml) by using butanol extracts of *A. absinthium*. On the contrary, *Listeria monocytogenes* showed the highest value (38.14±0.01 mg/ml) while *Staphylococcus aureus* showed the lowest one (81.43±0.01 mg/ml) in chloroform extract treatment. On the other side, methanol extract exhibited MIC value with 31.78±0.01 mg/ml on *Listeria monocytogenes*. In *A. herba alba* extracts, *Salmonella enterica* showed the same highest MIC value in both susceptible organic extracts 285.10±0.01 and mg/ml while *Listeria monocytogenes* showed the lowest MIC values 31.78±0.01 and 6.36±0.01 mg/ml in butanol and diethyl ether extracts respectively.

MBC values were ranged from 1.03±0.01 to 8.0±0.01 mg/ml in butanol and chloroform extracts of *A. absinthium* but *Listeria monocytogenes* showed MBC value 24.01±0.01 mg/ml in chloroform extract. However, they were ranged from 0.32±0.25 to 4.0±0.01 mg/ml of *A. herba alba* but the same pathogenic bacteria showed 20.0±0.01 mg/ml in butanol extract.

### 2. Phyto-characters analysis

43 morphological characters were obtained from both studied plant species. They are classified into 34 qualitative and 9 quantitative ones in 7 groups; Habitat (7 traits), Leaf (10 traits), Stem (4 traits), Petiole (1 trait), Bract (4 traits), Inflorescence (13 traits), Cypsela (4 traits) (**Table 10**) Similarity, there were 66 anatomical characters of both plant species (48 qualitative and 18 quantitative) which were distributed in 5 groups; Leaf (13 traits), Stem (17 traits), Root (8 traits), Leaf epidermis (8 traits), Stomata (20 traits) **(Tables 11 & 12)**. Moreover, Palynological characters were 25 as 12 qualitative and 13 quantitative. Each plant trait was symbolized, *i.e.* (+) present, (-) absent. **(Table 13) (Fig. 8-11).** Similarity character percentage was measured for each plant group character as shown in **(Table 14**).

**Fig. 8.**
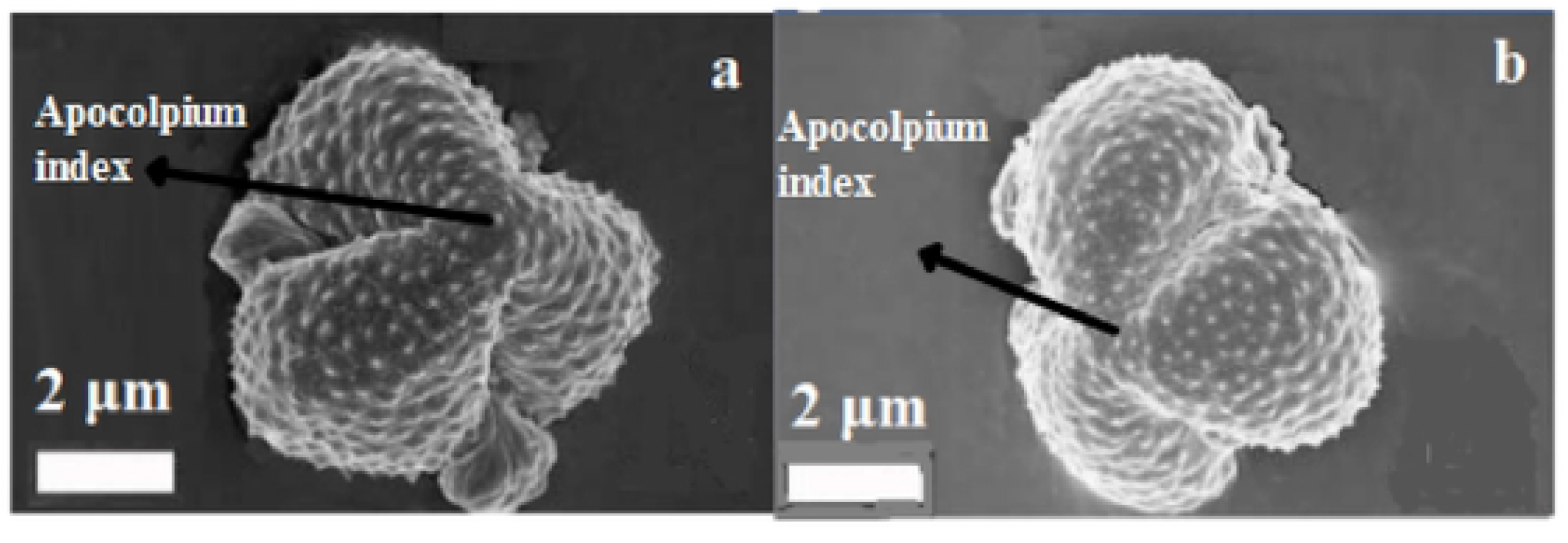
Apocolpium indexes of pollen grains for a. *A. absinthium* and b. *A. herba alba*.

**Fig. 9.**
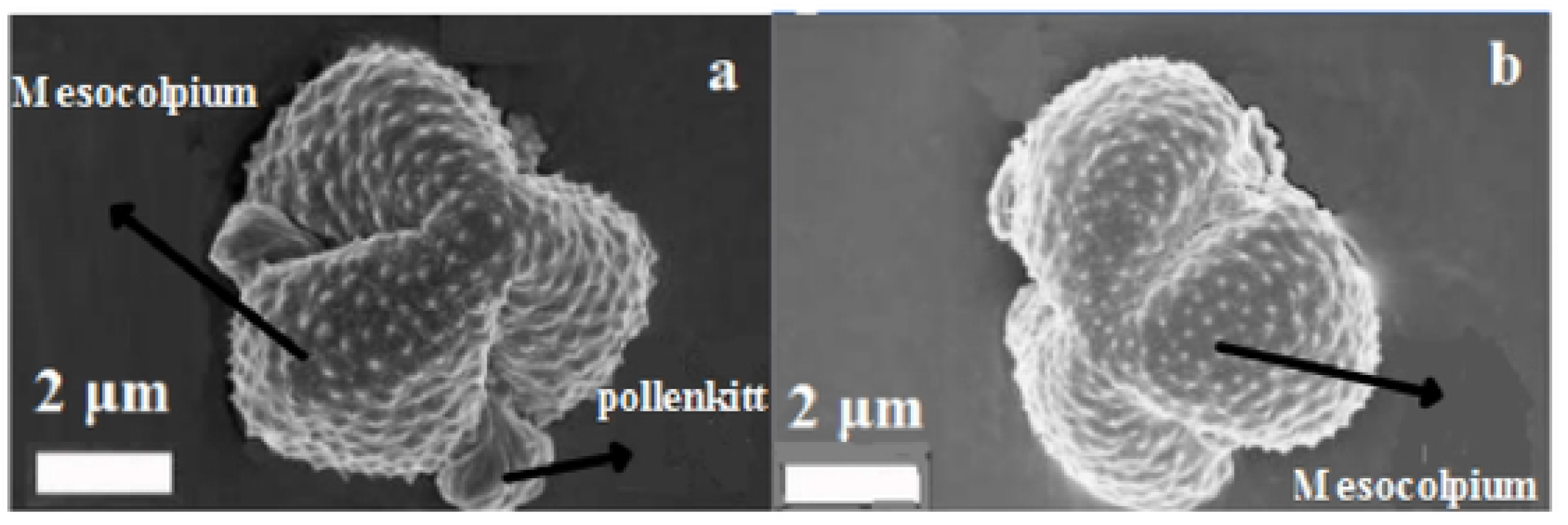
Mesocolpia of pollen grains for a. *A. absinthium* and b. *A. herba alba*.

**Fig. 10.**
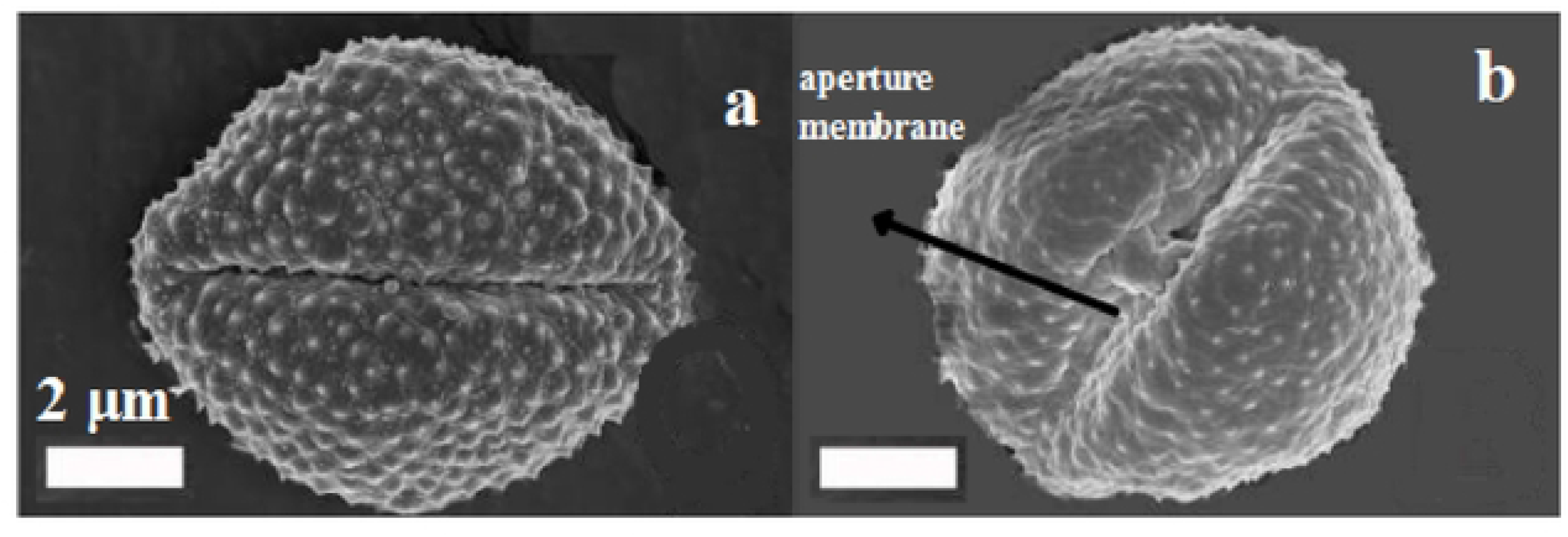
Apertures of pollen grains for a. *A. absinthium* and b. *A. herba alba*.

**Fig. 11.**
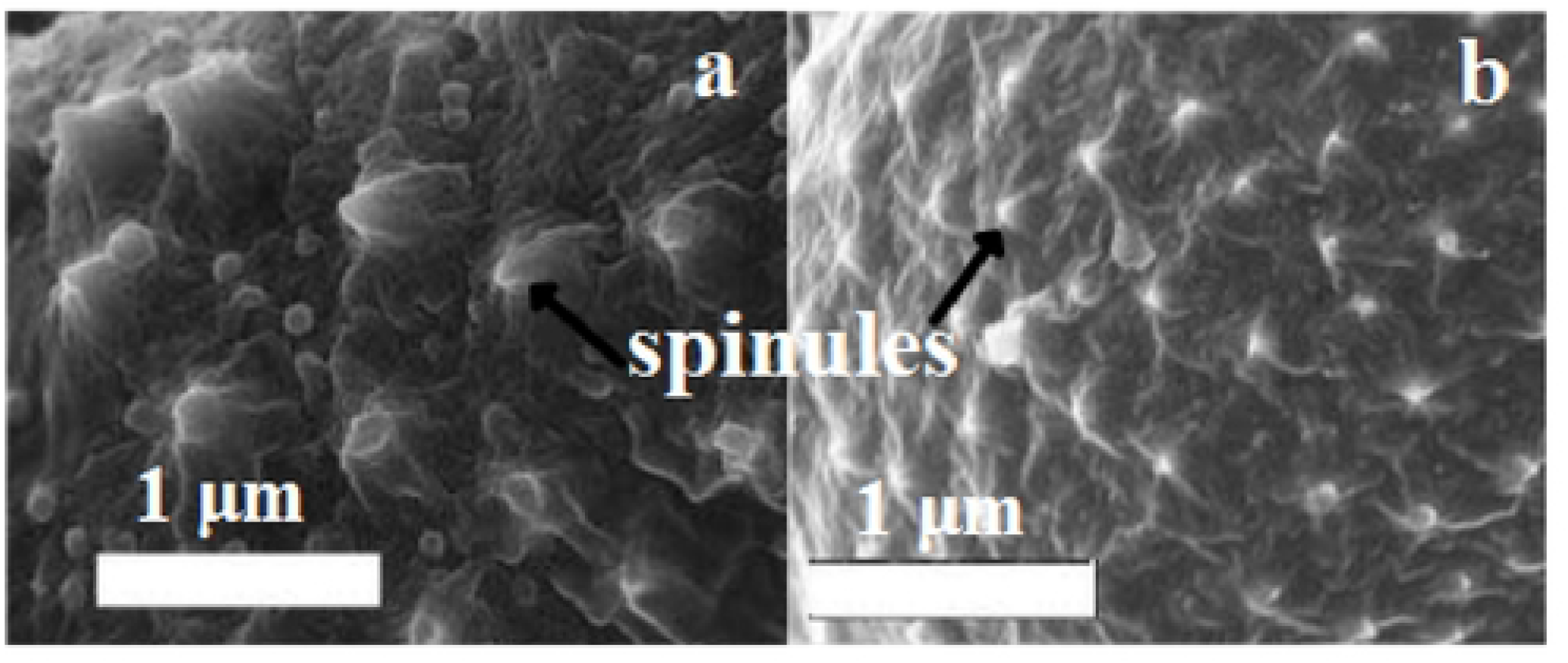
Spinules of pollen grains for a. *A. absinthium* and b. *A. herba alba*.

**Table 10.** Morphological characters of the studied two plant species.

| Morphological Characters | *A. absinthium* | *A. herba alba* |
| --- | --- | --- |
| habit | perennial(+) | perennial(+) |
| habit | herb(+) | herb(+) |
| tall | 70 to 150 cm(+) | 25 to 40 cm(-) |
| branching | numerous branches(+) | numerous branches(+) |
| plant not thistle-like | + | + |
| plant without latex | + | + |
| Pappus of bright setae | + | + |
| leaf blade | ovate to elliptic or ovate(+) | ovate to elliptic or ovate(+) |
| leaf dimension | 9-12 cm X 7-9 cm(+) | 0.2-0.3 cm X 0.1-0.2 cm(-) |
| leaf type | 3-pinnatisect(+) | 3-pinnatisect(+) |
| lobules | lanceolate-elliptic or linear(+) | lanceolate-elliptic or linear(+) |
| lobules dimension | 7-15 mm X 2-5 mm(+) | 2-3 mm X 0.5-1 mm(-) |
| lobule apex | obtuse apex(+) | obtuse apex(+) |
| leaves alternate | + | + |
| adaxial leaf surface | grayish-green(+) | dark green(-) |
| abaxial leaf surface | silver-gray(+) | clear green(-) |
| petiole | 2 to 6 cm long(+) | sessile(-) |
| stem hair | pubescence appressed(+) | pubescence appressed(+) |
| stem position | erect(+) | erect(+) |
| stem color | greenish gray(+) | greenish gray(+) |
| leafy bracts | 3-lobed(+) | bracts imbricate, multiseriate(-) |
| leafy bracts | entire(+) | entire(+) |
| leafy bracts | sessile(+) | sessile(+) |
| leafy bracts | oblong(+) | oblong(+) |
| bracts not spiny tipped | + | + |
| spherical baskets | 2.5-4 mm in diameter(+) | <3 mm(-) |
| florets | 2-4 in number(+) | 2-4 in number(+) |
| outer tubular flowers | pistillate(+) | pistillate(+) |
| inner funnel flowers | bisexual(+) | bisexual(+) |
| flower color | yellow(+) | yellow(+) |
| corolla | 1-2 mm(+) | 0.5-0.75 mm(-) |
| corolla | glandular(+) | glandular(+) |
| capitula | Globose(+) | Oblong(-) |
| capitula | 3.5 to 4 mm in diameter(+) | 0.25 to 0.3 mm(-) |
| capitula | oblong(+) | oblong(+) |
| receptacle | convex, semiglobose(+) | convex, semiglobose(+) |
| receptacle | covered with white ribbon-like scaly films(+) | naked(-) |
| receptacle | densely hairy(+) | absent(-) |
| involucre 2-seriate | + | + |
| heads homogamous | + | + |
| cypselas | oblong(+) | oblong(+) |
| cypselas | minute upper crown(+) | minute upper crown(+) |
| cypselas | mostly <0.5 mm(+) | mostly <0.5 mm(+) |
| cypselas | glabrous(+) | glabrous(+) |

**Table 11.** Anatomical characters of the studied two plant species (Root and Stem).

| Root and Stem Anatomy Characters | *A. absinthium* | *A. herba alba* |
| --- | --- | --- |
| a multilayered exodermis in root | + | + |
| below the exodermis, a cortex is present in root | + | + |
| below cortex, secondary phloem and some groups of sclerenchyma fibers in root | + | + |
| the secondary xylem is the dominant part of the root composed of vessels and tracheids | + | + |
| Endodermal secretory canals are present in the roots | + | - |
| root secretory canals, endodermal | + | - |
| root secretory canals, nonendodermal cortex | + | - |
| root secretory canals, secondary phloem | - | - |
| irregular pentagonal shape in young stem | + | + |
| more or less round or polygonal in old stem | + | + |
| one-layered epidermis, composed of oval to isodiametric cells of stem | + | + |
| prominent ribs in young stems contained collenchyma ribs in young stems contained collenchyma tissue | + | + |
| chlorenchyma is present between the ribs in stem | + | + |
| the periderm is continuous in older stems and consisted of a several layers of enlarged cells arranged in radial rows | + | + |
| the vascular bundles are collateral and arranged in a circle. The vascular bundles are collateral and arranged in a circle separated from one another by a parenchyma tissue in stem | + | + |
| the primary xylem consists of four to eight parallel rows of xylem elements; each row comprised 2–5 vessels. The vascular cylinder in a secondary state of growth produces secondary xylem inside and secondary phloem outside, giving to more basal parts of the stem almost cylindrical outline, while medullary rays on xylem side form connective tissue of lignified cells in stem | + | + |
| well-lignified sclerenchyma is above the phloem in stem | + | + |
| a large parenchyma cells are in the central region of the stem | + | + |
| small secretory canals in the cortex of stem | + | + |
| small secretory canals in the pith in stem | + | - |
| numerous nonglandular as well as glandular trichomes on stems | + | + |
| the non glandular trichomes are T-shaped, with various variable number of cells which form a neck of the trichome, and with long curly or straight arms in stem surface | + | + |
| glandular trichomes are of biseriate type covered with cuticle sheath in stem | + | + |
| stem Secretory canals, cortex | + | + |
| stem Secretory canals, pith | + | - |

**Table 12.**
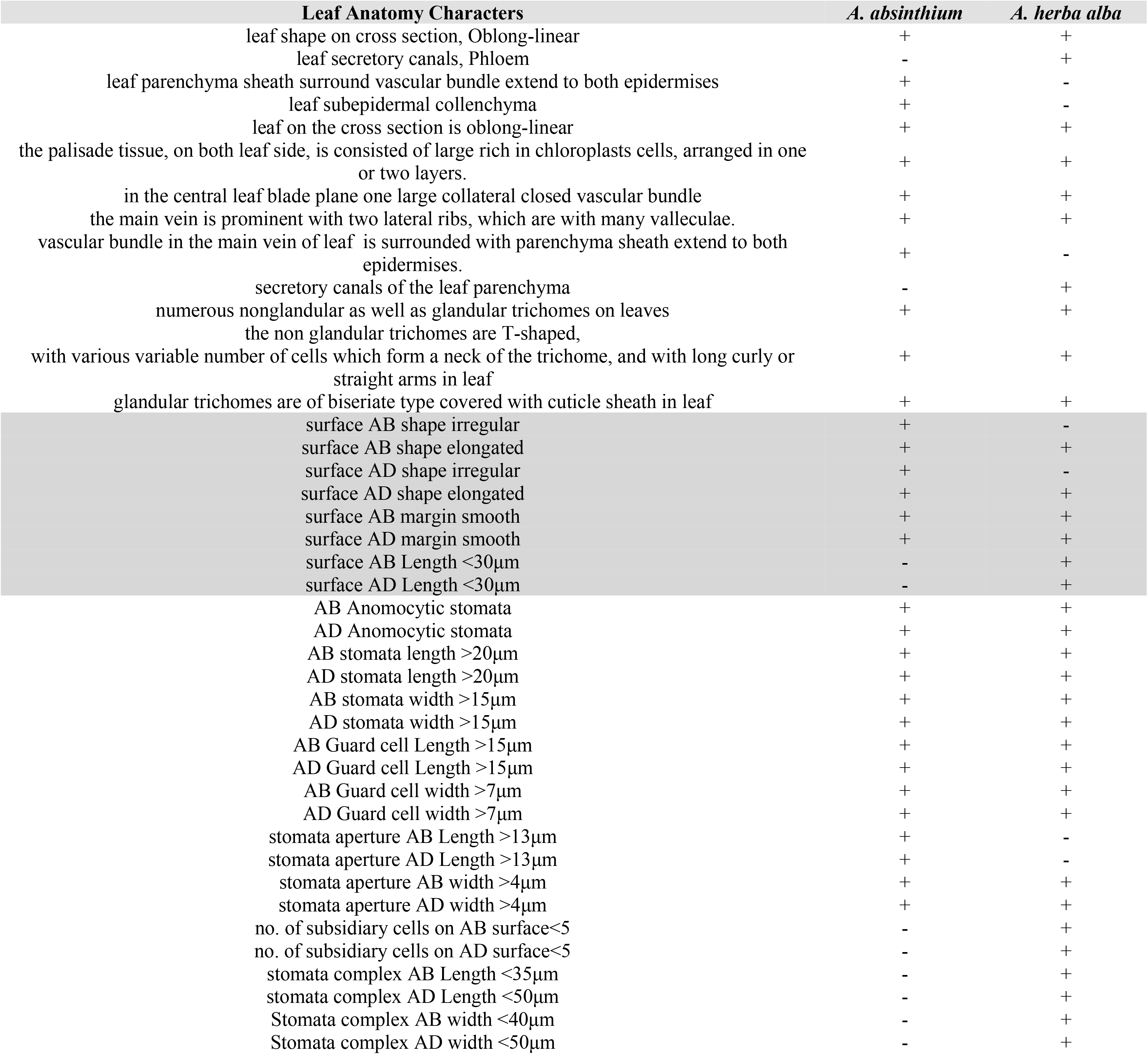
Anatomical characters of the studied two plant species (Leaf, Leaf epidermis and Stomata).

**Table 13.**
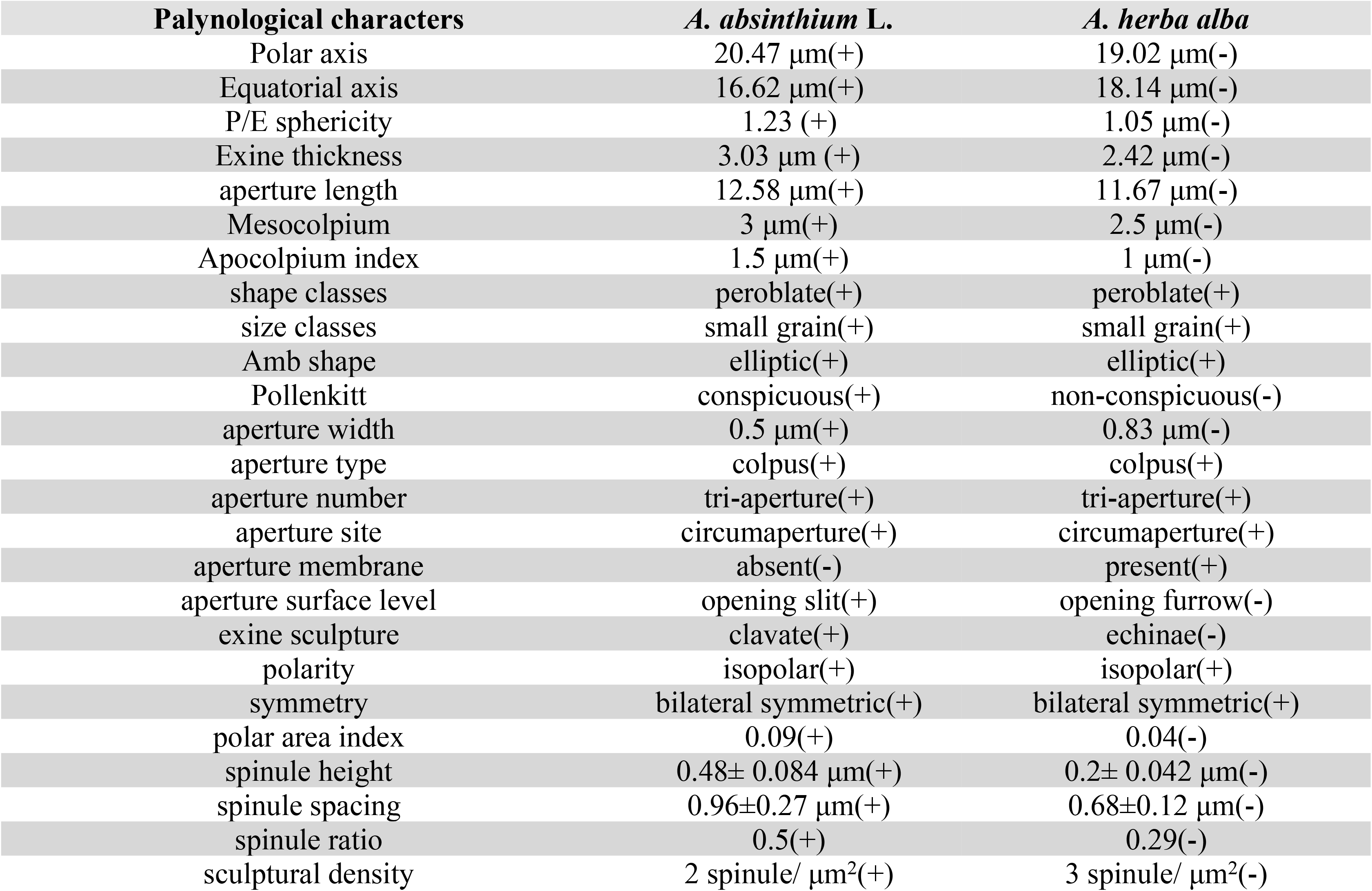
Palynological characters of the studied two plant species.

| Palynological characters | *A. absinthium* L. | *A. herba alba* |
| --- | --- | --- |
| Polar axis | 20.47 $\mu\text{m}$(+) | 19.02 $\mu\text{m}$(-) |
| Equatorial axis | 16.62 $\mu\text{m}$(+) | 18.14 $\mu\text{m}$(-) |
| P/E sphericity | 1.23 (+) | 1.05 $\mu\text{m}$(-) |
| Exine thickness | 3.03 $\mu\text{m}$ (+) | 2.42 $\mu\text{m}$(-) |
| aperture length | 12.58 $\mu\text{m}$(+) | 11.67 $\mu\text{m}$(-) |
| Mesocolpium | 3 $\mu\text{m}$(+) | 2.5 $\mu\text{m}$(-) |
| Apocolpium index | 1.5 $\mu\text{m}$(+) | 1 $\mu\text{m}$(-) |
| shape classes | peroblate(+) | peroblate(+) |
| size classes | small grain(+) | small grain(+) |
| Amb shape | elliptic(+) | elliptic(+) |
| Pollenkitt | conspicuous(+) | non-conspicuous(-) |
| aperture width | 0.5 $\mu\text{m}$(+) | 0.83 $\mu\text{m}$(-) |
| aperture type | colpus(+) | colpus(+) |
| aperture number | tri-aperture(+) | tri-aperture(+) |
| aperture site | circumaperture(+) | circumaperture(+) |
| aperture membrane | absent(-) | present(+) |
| aperture surface level | opening slit(+) | opening furrow(-) |
| exine sculpture | clavate(+) | echinae(-) |
| polarity | isopolar(+) | isopolar(+) |
| symmetry | bilateral symmetric(+) | bilateral symmetric(+) |
| polar area index | 0.09(+) | 0.04(-) |
| spinule height | 0.48± 0.084 $\mu\text{m}$(+) | 0.2± 0.042 $\mu\text{m}$(-) |
| spinule spacing | 0.96±0.27 $\mu\text{m}$(+) | 0.68±0.12 $\mu\text{m}$(-) |
| spinule ratio | 0.5(+) | 0.29(-) |
| sculptural density | 2 spinule/ $\mu\text{m}^2$(+) | 3 spinule/ $\mu\text{m}^2$(-) |

**Table 14.**
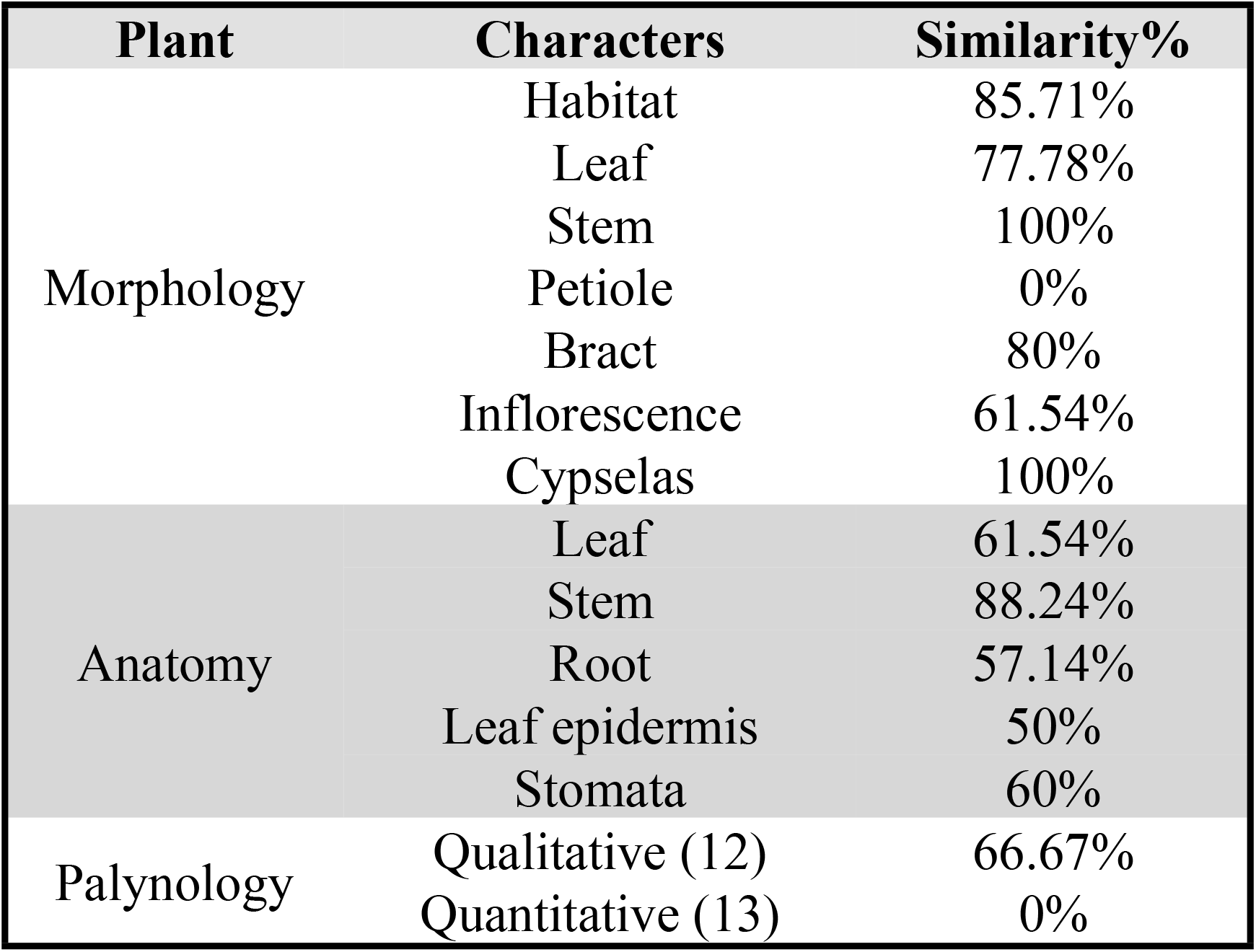
Similarity character percentage of two studied plant species

| Plant | Characters | Similarity% |
| --- | --- | --- |
| Morphology | Habitat | 85.71% |
|  | Leaf | 77.78% |
|  | Stem | 100% |
|  | Petiole | 0% |
|  | Bract | 80% |
|  | Inflorescence | 61.54% |
|  | Cypsels | 100% |
| Anatomy | Leaf | 61.54% |
|  | Stem | 88.24% |
|  | Root | 57.14% |
|  | Leaf epidermis | 50% |
|  | Stomata | 60% |
| Palynology | Qualitative (12) | 66.67% |
|  | Quantitative (13) | 0% |

### 3. Statistical Analysis

The analysis of variance for various MIC values by ANOVA test showed significant differences for all tested bacterial strains. In *A. absinthium* extracts, *Listeria monocytogenes* versus *Pseudomonas aeruginosa* showed high significant with *F* test equals 51.01 while *Pseudomonas aeruginosa* versus *Salmonella enterica* showed low significant with *F* test equals 1.64. In *A. herba alba* extracts, *Listeria monocytogenes* versus *Salmonella enterica* showed high significant with *F* test equals 111.61 while *Pseudomonas aeruginosa* versus *Salmonella enterica* showed low significant with *F* test equals 2.57. Pearson correlation coefficients among MICs of *A. absinthium* and *A. herba alba* extracts against the tested bacterial strains showed that *Pseudomonas aeruginosa* had a highly positive correlation, *Salmonella enterica* and *Staphylococcus aureus* had moderate correlations and *Listeria monocytogenes* had low correlation. Regression describes the co-variation among MIC variables. SLR curve indicates the significant relationships among them. It was extremely high regressed in *Pseudomonas aeruginosa*, high regressed in both *Salmonella enterica* and *Staphylococcus aureus* and moderate regressed in *Listeria monocytogenes*. Finally, all comparative data between two studied plant species differentiating into antibacterial characters, morphological, anatomical, palynological traits was represented by simple linear regression curve into high regressed (Table 15) (**Fig. 12 & 13)**.

**Fig. 12.**
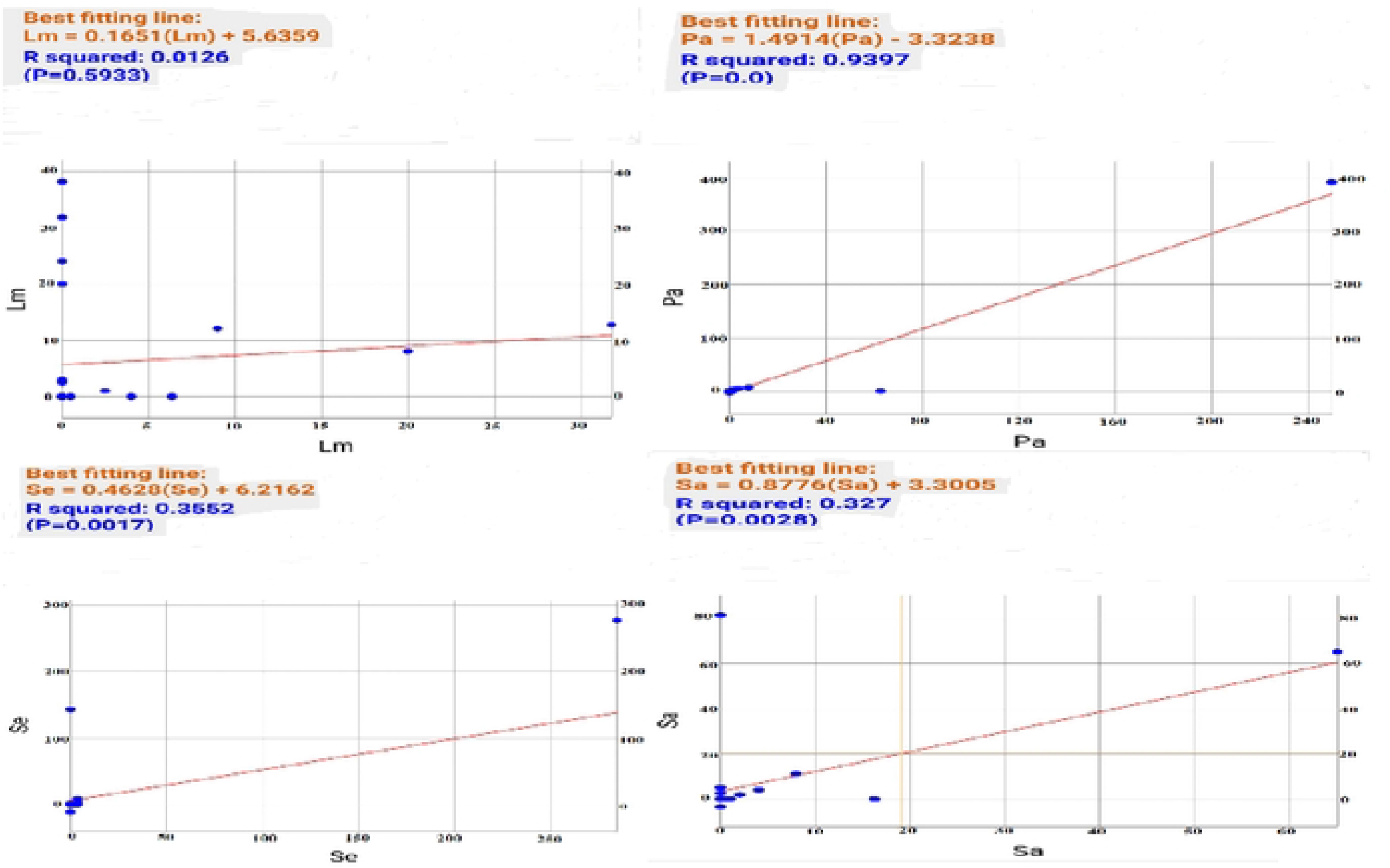
SLR curves of MICs of both plant species extracts against the tested bacterial strains

**Fig. 13.**
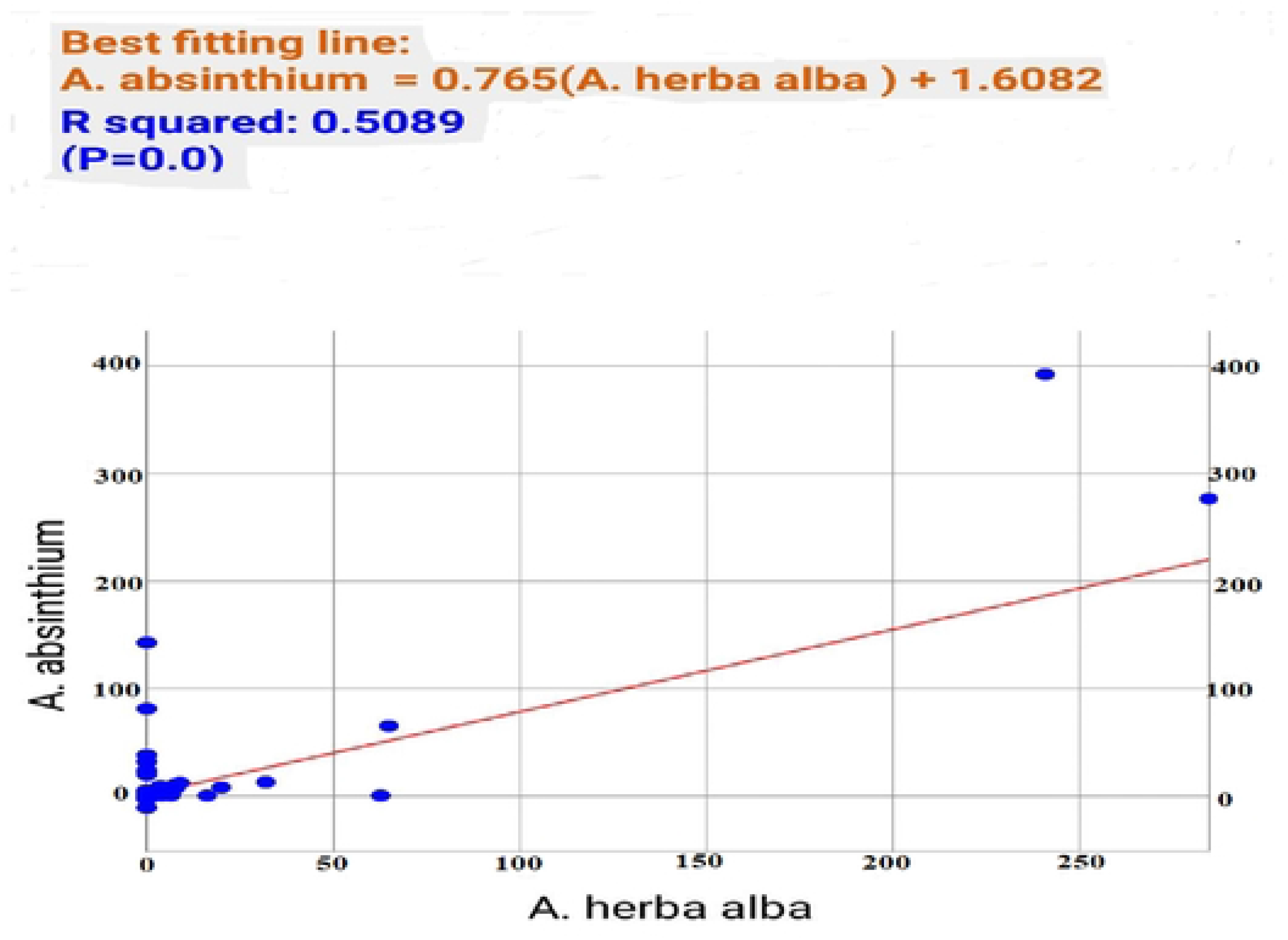
SLR curve for all comparative data of both plant species.

**Table 15.** Pearson correlation coefficients among MICs of both plant species extracts against the tested bacterial strains.

|  | *Listeria monocytogenes* | *Pseudomonas aeruginosa* | *Salmonella enterica* | *Staphylococcus aureus* |
| --- | --- | --- | --- | --- |
| *Listeria monocytogenes* | 0.112 |  |  |  |
| *Pseudomonas aeruginosa* |  | 0.969 |  |  |
| *Salmonella enterica* |  |  | 0.596 |  |
| *Staphylococcus aureus* |  |  |  | 0.571 |

## Discussion

The searching for medicinal plant extracts exhibiting antimicrobial activity is increased due to the WHO reports indicating the rise of antimicrobial resistance **(Kebede *et al*., 2021)**. The choice of organic solvent is the proper use in extraction of natural resources. It was clear in this study which covered many number of different organic solvents besides aqueous one. The butanol solvent was the most convenient extract in this investigation due to moderate polarity index (4) with low solubility in water (0.43%) comparing to other organic solvents. Chloroform ranked the best second organic solvent due to less viscosity (0.57 cP). The third effective extract was diethyl ether due to less in both of polarity index (2.8) and viscosity (0.32 cP) with high solubility in water (6.89%) **(Paul 2002)**.

According to MICs values, *Listeria monocytogenes* was the most susceptible with less MIC value while *Pseudomonas aeruginosa* was the most resistant with high MIC value of *A. absinthium* by using butanol extract. On the other hand, *Listeria monocytogenes* was the most susceptible with less MIC value of *A. herba alba* by using diethyl ether extract while *Salmonella enterica* was the most resistant with high MIC value of *A. herba alba* by using both investigated organic extracts. It may refer to the structure of cell wall and its interaction with extracted chemical components with different affinity in membrane permeability and potentiality of protein binding because *Listeria monocytogenes* is Gram-positive while others are Gram-negative (**Mariola *et al*., 2022)**.

For MBCs values, *Listeria monocytogenes* needed high inoculum dose to propagate in *A. absinthium* and *A. herba alba* treatments for chloroform and butanol extracts respectively whereas *Pseudomonas aeruginosa* needed the lowest inoculum dose in all treatments that may refer to be more virulent than the preceding.

**Maria *et al*. 2023** demonstrated the antibacterial activity of *A. absinthium* by using ethanol extract against *Listeria monocytogenes* and *Staphylococcus aureus*. This plant material collected from Europe especially the outskirts of Blaj, Alba, Romania. **Marija *et al*. 2021** confirmed last evidence with other studied bacterial strains; *Pseudomonas aeruginosa* and *Salmonella enterica* collecting from also Europe especially among Serbian population. **Roman *et al*., 2021** reported on negative inhibitory effect on *Staphylococcus aureus* by using 40% or 70% ethanol extract of *A. absinthium* and positive inhibitory one by using 90% ethanol extract. From previous studies, it is clear that the chemical constituent of *A. absinthium* differs due to the physiological parts **(Riahi *et al*., 2013)**, geographical areas **(Msaada, *et al*., 2015)**, the degree of senescence **(Lommen *et al*., 2006)** and temperature **(Nguyen, *et al*., 2018)**. It implies on antimicrobial activities of such plant material with variety of degrees according to different geographical regions (**Maria *et al*. 2023)**.

Studies on micromorphological features besides traditional ones have gained more attention nowadays in plant taxonomy **(Srilakshmi and Naidu, 2014)**. There are further studies on *Artemisia* sp. which could support or contradict the present evidences and nuances among them. Nevertheless, the full gathering data is not sufficient to resolve taxonomic issues within the genus *Artemisia.* **(Javad *et al*., 2022; Li-Li *et al*., 2022)**. This study gives obvious picture of all descriptive data for both plant species and try to solve taxonomist’s perceptions in the delimitation of *Artemisia* species by paving the way to classify them comparatively with other *Artemisia* species and be useful for subgeneric classification of the genus *Artemisia*. *P*-values indicated that the two studied plant species were effective for this investigation. They reflected the importance of their natural ingredients which can be used as natural alternatives against microbial infections. Pearson correlation coefficients confirmed that the both plant species were the best inhibitory agent against *Pseudomonas aeruginosa*. Furthermore, (SLR) equations recognized the data analysis as exponential curves revealing that the both plant species were compatible between them from one side and among bacterial strains from another side.

Based on our research and the aforementioned studies, we can state that the application of new therapeutic protocols against resistant infectious pathogenic diseases on the fundamental of *Artemisia* extracts is a real possibility that should be taken into considerations.

## Conclusion

1- The need for natural alternatives becomes the world demand nowadays
2- *Artemisia absinthium* and *A. herba alba* have the best antimicrobial activities
3- Butanol solvent records the most favorable natural extract.
4. Geographical region influences on the active chemical composition of plant extract.
5. Micro-botanical characters enhance traditional ones in taxonomical approaches.

## Abbreviations

SEM: Scanning Electron Microscope
STL: Simple Linear Regression
WHO: World Health Organization

### Glossary

**Amb (Erdtman, 1952)**: Outline of pollen grain viewed with one of the poles exactly uppermost. **Aperture (Erdtman, 1947)**: A weak, performed opening of the exine that permits the exit of intra-exinous contents(e.g. pollen tube).

**Apocolpium index (Erdtman, 1952)**: Area at a pole, delimited towards the equator by the polar limits of the mesocolpia.

**Circumaperturate (Straka, 1964)**: Describing a pollen grain with equatorial apertures that are regularly arranged around a circular outline.

**clavate (Proctor and Yeo, 1973):** spinules of sharp point apex with wide beneath on pollen grain surface.

**Colpus (Erdtman, 1943)**: Longitudinal aperture having a length greater than twice its width. **Echinae (Proctor and Yeo, 1973)**: spinules of pointed apex with narrow beneath on pollen grain surface.

**Equatorial axis (Erdtman, 1943)**: A line, lying in the equatorial plane, perpendicular to the polar axis and passing through it.

**Exine (Fritzsche, 1837)**: The main, outer, usually resistant layer of a sporoderm.

**Mesocolpium (Erdtman, 1952)**: The area of the grain surface between two adjacent colpi. It is usually delimited transverse lines drawn through the polar ends of the colpi.

**P/E sphericity (Erdtman, 1943)**: The ratio of the length of the polar axis (P) to the equatorial axis (E).

**Peroblate (Erdtman, 1943)**: Describing the shape of a pollen grain or spore in which the ratio between the polar axis and the equatorial axis is less than 2.

**Polar area index (Iversen and Troels-Smith, 1950)**: the ratio of the distance between the apices of two colpi to its equatorial axis.

**Polar axis (Wodehouse, 1935)**: A straight line connecting the distal and proximal pole of the pollen grain.

**Pollenkitt (Knoll, 1930)**: A sticky material, produced by the tapetum, that may hold pollen grains together during dispersal.

**Sculpture (Kuprianova, 1948)**: The surface relief, or topography, of a pollen grain.

**Sculpural density (Balme, 1988)**: The estimated number of sculptural elements in an area of 100 μm2 of the surface of the exine.

**spinules (Proctor and Yeo, 1973)**: processes are protruded from exine sculpture.

## Data Availability

All datasets were achieved, used and analyzed in the current study. They were included within this manuscript.

## Declaration

The authors pronounced that this suggested work is not financially supported from any other financier.

## Authors’ Contributions

Abdullah Mashraqi, Mohamed A. Al Abboud, Khatib Sayeed Ismail, Yosra Modafer, Mukul Sharma and A. El-Shabasy contributed equally to this work.

## Acknowledgments

The authors extend their appreciation to the Deputyship for Research & Innovation, Ministry of Education in Saudi Arabia for funding this research work through project number ISP22-24.

